# Non-competitive resource exploitation within-mosquito shapes evolution of malaria virulence

**DOI:** 10.1101/149443

**Authors:** G. Costa, M. Gildenhard, M. Eldering, R.L. Lindquist, A.E. Hauser, R. Sauerwein, C. Goosmann, V. Brinkmann, E.A. Levashina

**Affiliations:** Max Planck Institute for Infection Biology, Berlin, Germany; Radboud University Medical Center, Nijmegen, The Netherlands; German Rheumatism Research Centre (DRFZ), Berlin, Germany; Charité-Universitätsmedizin Berlin, Germany

**Author notes:** These authors contributed equally to this work (M. Gildenhard, M. Eldering).

## Abstract

Malaria is a fatal human parasitic disease transmitted by a mosquito vector. The evolution of within-host malaria virulence has been the focus of many empirical and theoretical studies. However, the vector’s contribution to virulence evolution is not well understood. Here we explored how within-vector resource exploitation impacts evolutionary trajectories of within-host *Plasmodium* virulence. We developed a nested model of within-vector dynamics and malaria epidemiology, which predicted that non-competitive resource exploitation within-vector restricts within-host parasite virulence. To validate our model, we experimentally manipulated mosquito lipid trafficking and gauged within-vector parasite development, within-host infectivity and virulence. We found that mosquito-derived lipids determine within-host parasite virulence by shaping development and metabolic activity of transmissible sporozoites. Our findings uncover the role of within-vector environment in regulating within-host *Plasmodium* virulence and identify *Plasmodium* metabolic traits that may contribute to the evolution of malaria virulence.

## INTRODUCTION

Malaria is caused by the vector-borne protozoan parasite *Plasmodium falciparum* and kills 429,000 people annually, predominantly in sub-Saharan Africa^1^. The unacceptably high malaria mortality is tightly linked to within-host parasite virulence (here defined as capacity to cause harm to the human host). Therefore, predicting the evolution of *Plasmodium* virulence has important epidemiological, socio-medical and evolutionary implications for designing malaria control strategies. The life cycle of malaria parasites is split between the host and the mosquito vector. Theoretical and empirical studies focused on regulation of within-host malaria virulence, including contributions of parasite genetic factors and host immune responses, and on the link between virulence and host-to-vector transmission^2–4^. The latter reports showed that genetically-encoded within-host parasite virulence increases the production of infective to vector forms and, thereby, parasite transmission to the vector^5^. Females of anopheline mosquitoes feed on blood to initiate ovary development. Therefore, only females ingest *Plasmodium* and contribute to malaria transmission. During a blood meal, blood-borne sexual forms of parasites fuse to form motile zygotes that within two days after infection traverse the midgut epithelium, reach the basal side of the midgut to round-up into oocysts. In the next 10-15 days, each oocyst undergoes sporogony, a process that generates thousands sporozoites, the parasite stage infective to humans. Sporozoites egress from the oocysts and migrate to the mosquito salivary glands to reach full maturity and infectivity^6^. Nutrients taken up by female mosquitoes during a blood meal are essential for vector reproduction and for massive *Plasmodium* proliferation at the oocyst stage^7,8^. Although many studies examined associations between *Plasmodium* virulence and vector fitness, only a few reported significant changes in longevity and fecundity of *Plasmodium*-infected mosquitoes, thus precluding generalized conclusions^9–13^. Whether within-vector resource exploitation by parasites modulates within-host malaria virulence has never been explored.

Here we applied theoretical and experimental approaches to examine the link between parasite within-vector resource exploitation and virulence towards the mammalian host. We modeled alternative scenarios of vector-parasite symbiosis and investigated their consequences for parasite transmission dynamics. Our modeling results predicted that vector-parasite relationship of resource acquisition shapes the evolution of within-host virulence. A parasitic relationship, where the parasite exploited vector mechanisms to scavenge resources, restricted parasite virulence, whereas virulence runaway was observed in a competitive relationship. To test these predictions, we experimentally manipulated lipid trafficking in genetically identical mosquitoes and exploited a rodent malaria model to gauge the impact of these manipulations on within-host virulence of genetically identical parasites. We found that mosquito-derived lipids determine within-host *Plasmodium* virulence by shaping sporogony and metabolic activity of sporozoites. By showing that evolution of malaria virulence is metabolically regulated by the within-vector nutritional environment, our findings have important implications for vector control strategies based on biological competitors or on manipulation of mosquito reproduction.

## RESULTS

### Within-vector environment shapes evolution of within-host malaria virulence

To test for the evolutionary consequences of a trade-off between mosquito reproduction, parasite reproduction and virulence, we used a modeling approach by considering a within-vector dynamics model in an epidemiological framework (Materials and Methods). We modeled within-vector resource competition between mosquito reproduction (egg production) and parasite reproduction (sporozoite load) as a competitive Lotka-Volterra equation^14,15^. This model describes egg production, parasite production and the competition strength between vector and parasite *α*_*EP*_ and *α*_*PE*_ (their ability to inhibit each other’s growth). Mechanistically, the parasite’s competition coefficient represents its within-vector resource scavenging and, thereby, its harm towards vector reproduction.

We next combined this within-vector model with an epidemiological model for vector-borne disease, which describes human-to-vector and vector-to-human parasite transmission. Pathogen evolution can be investigated with epidemiological models, assuming that selection maximizes the parasite’s ability to spread from one host to the next expressed as the basic reproductive ratio R_0_. For vector-borne diseases, R_0_ is the product of the vector-to-host and host-to-vector transmission potentials^16^. In line with previous reports^17–19^, we postulate that virulence towards the host is influenced by within-vector resource acquisition. We specifically hypothesize that within-vector resource exploitation influences virulence by impacting parasite load and infectivity to the host (Figure 1A).

**Figure 1.**
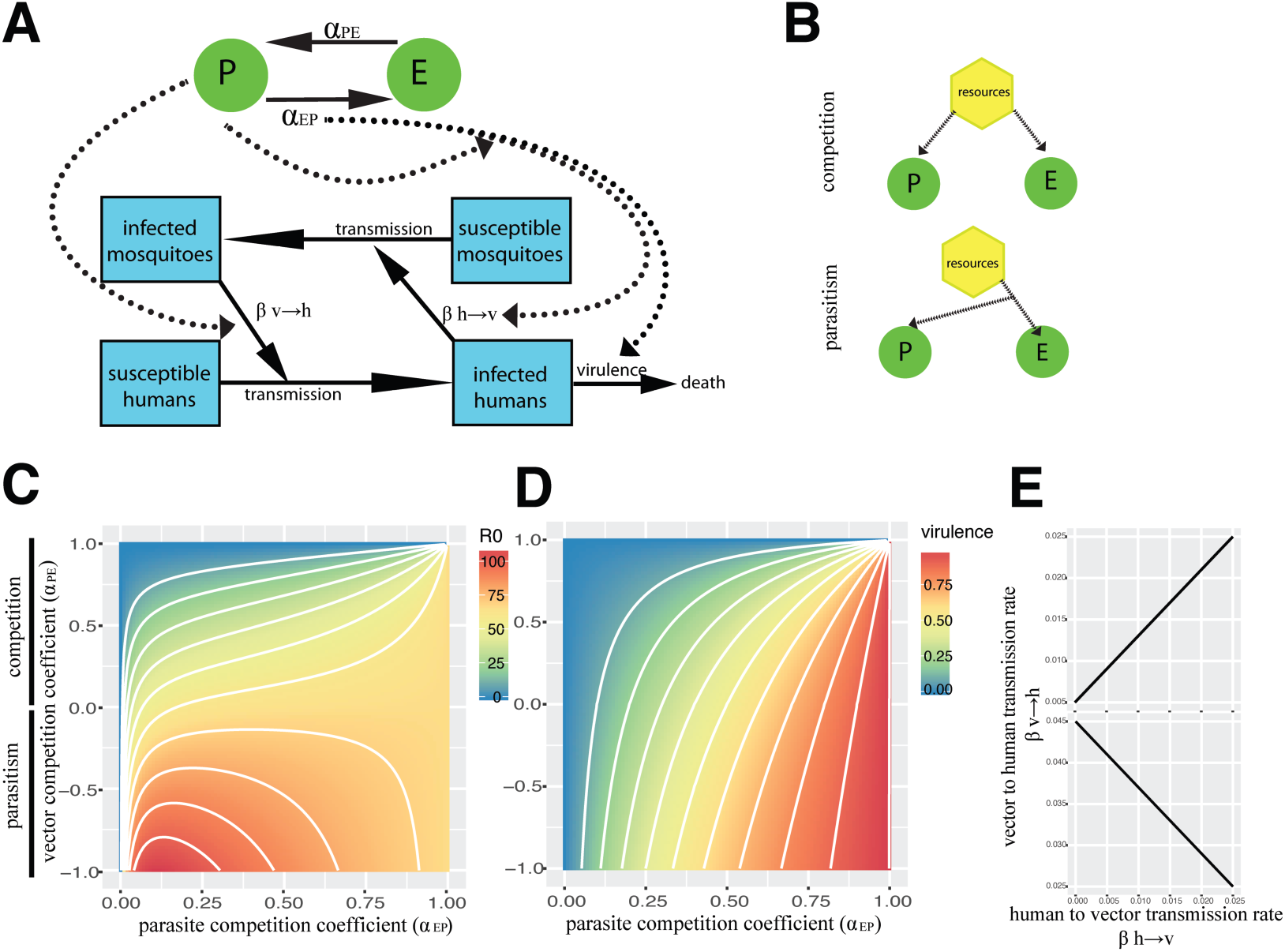
Evolution of vector-parasite antagonism and virulence. **(A)** Model scheme of parasite-vector non-mutualistic symbiotic relationship (green), nested into epidemiological dynamics (blue): within-vector parasite density is positively linked to vector-to-human transmission, whereas human-to-vector transmission is a linear function of the parasite’s virulence towards mosquito reproduction (competition coefficient) and human host longevity. **(B)** The biological relationship is competitive, if a common resource is scavenged, or parasitic if mosquito resource scavenging aids parasite development. **(C)** Contour plot of parasite fitness R_0_ under varying within-vector interaction terms. The y-axis shows the relationship of mosquito egg production towards the parasite *α*_*PE*_ over the parasite’s competition coefficient (i.e. virulence) *α*_*EP*_. Positive values of *α*_*PE*_ showcase a competitive parasite-vector interaction, whereas negative values imply that vector reproduction benefits parasite density leading to a parasitic relationship. **(D)** Contour plot of parasite virulence towards the host under varying within-vector interaction terms. Under competitive scenario, parasite fitness increases with virulence, whereas under parasitic scenario, an evolutionarily stable virulence strategy(ESV) emerges. **(E)** Vector-to-human transmission over human-to-vector transmission under varying degrees of parasite virulence. When mosquito and parasites independently scavenge and directly compete for resources (top, *α*_*PE*_ = 0.8), the transmission terms are positively linked. If parasites prey on mosquito resource scavenging (bottom, *α*_*PE*_ = −0.8), a trade-off between the transmission terms explains the emergence of the ESV. P = parasites; E = mosquito eggs.

Combining within-host Lotka-Volterra dynamics with epidemiological dynamics allows us to investigate evolution of R_0_ under varying parasite and vector competition coefficients. When vector and parasite use independent mechanisms to scavenge resources (competitive relationship), both competition coefficients have a positive sign (*α*_*EP*_ > 0; *α*_*PE*_ > 0), reflecting a reciprocally inhibitory competitive relationship (Figure 1B). We found that competitive environment selects for higher parasite’s competition coefficient (Figure 1C), eventually leading to vector sterilization and to high levels of within-host virulence (Figure 1D). This can be explained by the fact that both vector-to-human and human-to-vector transmission rates benefit from increasing the parasite’s competitiveness. When parasites mechanistically depend on the vector’s investment in reproduction (parasitic relationship), the vector’s competition coefficient turns negative (*α*_*EP*_ > 0; *α*_*PE*_ < 0) (Figure 1C). We found that in contrast to the competitive scenario, the parasitic relationship leads to opposing evolutionary trajectories between the two transmission events. Indeed, higher *α*_*EP*_ decreases parasite’s density and vector-to-human transmission but increases human-to-vector transmission (Figure 1E). Such a trade-off between transmission events would lead to the emergence of an evolutionarily stable malaria virulence (Figure 1C and D).

Taken together, our theoretical analyses predicted that the shape of vector-parasite relationship defines the evolutionary trajectories of within-host virulence.

### Mosquito lipid environment regulates *Plasmodium* virulence in the next mammalian host

To formalize our theoretical assumption that within-host virulence is defined by within-vector resource acquisition, we examined changes in *Plasmodium* virulence after restricting within-vector lipid resources. In these experiments, we used RNAi in adult females to deplete the major mosquito lipid transporter lipophorin (Lp) without affecting mosquito diet or development^20–22^. As expected, Lp depletion prevented lipid transport between mosquito tissues, inhibited ovary and parasite development (Figure S1). We next exposed naïve mice to the bites of *P. berghei*-infected control or Lp-depleted mosquitoes. We found that Lp depletion drastically attenuated parasite infectivity and virulence. Indeed, only 20% of mice bitten by Lp-depleted mosquitoes became infected and only 10% developed severe neurological symptoms of experimental cerebral malaria (ECM) compared to 90% in controls (Figure 2A-C). As the observed phenotypes could result from the differences in numbers of inoculated parasites, we subcutaneously injected naïve mice with equal quantities of sporozoites isolated from control and Lp-depleted mosquitoes. Again, fewer than 40% of mice injected with lipid-deprived sporozoites became infected and only 20% of mice developed ECM, as compared to 100% in controls (Figure 2D-F). These results confirm the major assumption of our model and show that the within-vector lipid environment impacts within-host parasite virulence.

**Figure 2.**
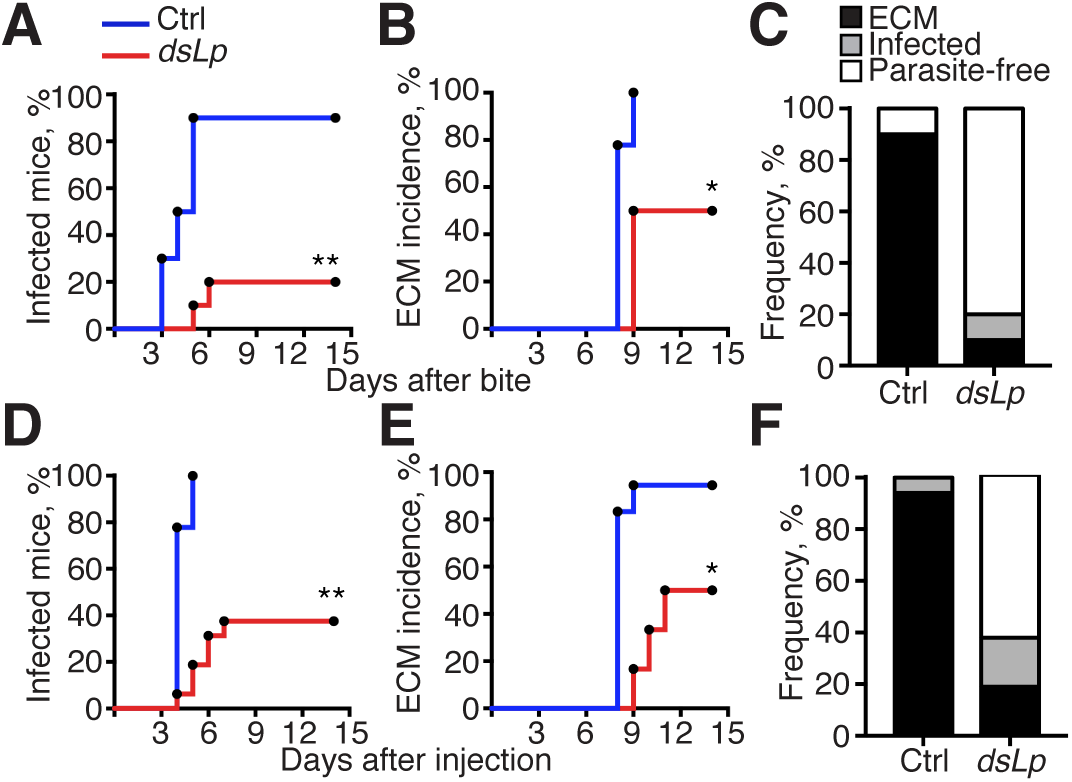
Within-vector environment impacts within-host *Plasmodium* virulence. C57BL/6 mice infected by bites of *P.* berghei-infected mosquitoes (**A-C**) or by sporozoite injection (**D-F**) were daily monitored for parasite occurrence in the blood and for symptoms of experimental cerebral malaria (ECM). (**A**) Kaplan-Meier analysis of time to infection (blood stage parasitaemia), (**B**) incidence (%) of experimental cerebral malaria (ECM) and (**C**) cumulative health status of mice infected by bites of control (Ctrl, n=10) or Lp-depleted (*dsLp*, n=10) mosquitoes. (**D**) Kaplan-Meier analysis of time to infection (blood stage parasitaemia), (**E**) incidence (%) of experimental cerebral malaria (ECM) and (**F**) cumulative health status of mice infected by subcutaneous injection of 5,000 sporozoites dissected from the salivary glands of control (Ctrl, n=18) or Lp-depleted (*dsLp*, n=16) mosquitoes. Asterisks indicate statistically significant differences (*: p<0.05;**: p<0.001; log-rank Mantel-Cox test and 2-way ANOVA).

### Plasmodium parasitizes mosquito lipids for productive sporogony and formation of infective sporozoites

Our theoretical study predicted that the type of symbiotic relationship (competitive or parasitic) defines the evolutionary trajectory of parasite virulence. As restricting trafficking of lipid resources impacted virulence, we examined the parasitic scenario, where *Plasmodium* exploits mosquito Lp for lipid delivery. We microscopically gauged lipid accumulation and parasite growth using Nile Red lipid staining, and observed accumulation of neutral lipids in the peripheral cytoplasm and vesicles of mature oocysts in control mosquitoes. In contrast, levels of Nile Red staining were significantly lower in the oocysts of Lp-depleted females (Figure 3A and B). Moreover, the immunofluorescence analysis with anti-Lp antibodies in controls showed an increase in the rate of Lp-positive oocysts from 25% to 72% at 7 and 13 days post infection (dpi), respectively (Figure S2). These results support the parasitic scenario, in which *Plasmodium* exploits Lp for lipid delivery to its proliferating stages. Depletion of Lp inhibited oocyst sporulation, caused abnormal cytoplasmic vacuolizations (Figure 3C and Figure S3), and significantly reduced the size of mature oocysts (Figure 3D). In line with these results, perturbation of lipid trafficking decreased the density of transmissible sporozoites (Figure 3E). Similar results were obtained with the human malaria parasite *P. falciparum* (Figure 3D and E), suggesting that within-vector parasitic relationships are conserved between human and rodent *Plasmodium* species.

**Figure 3.**
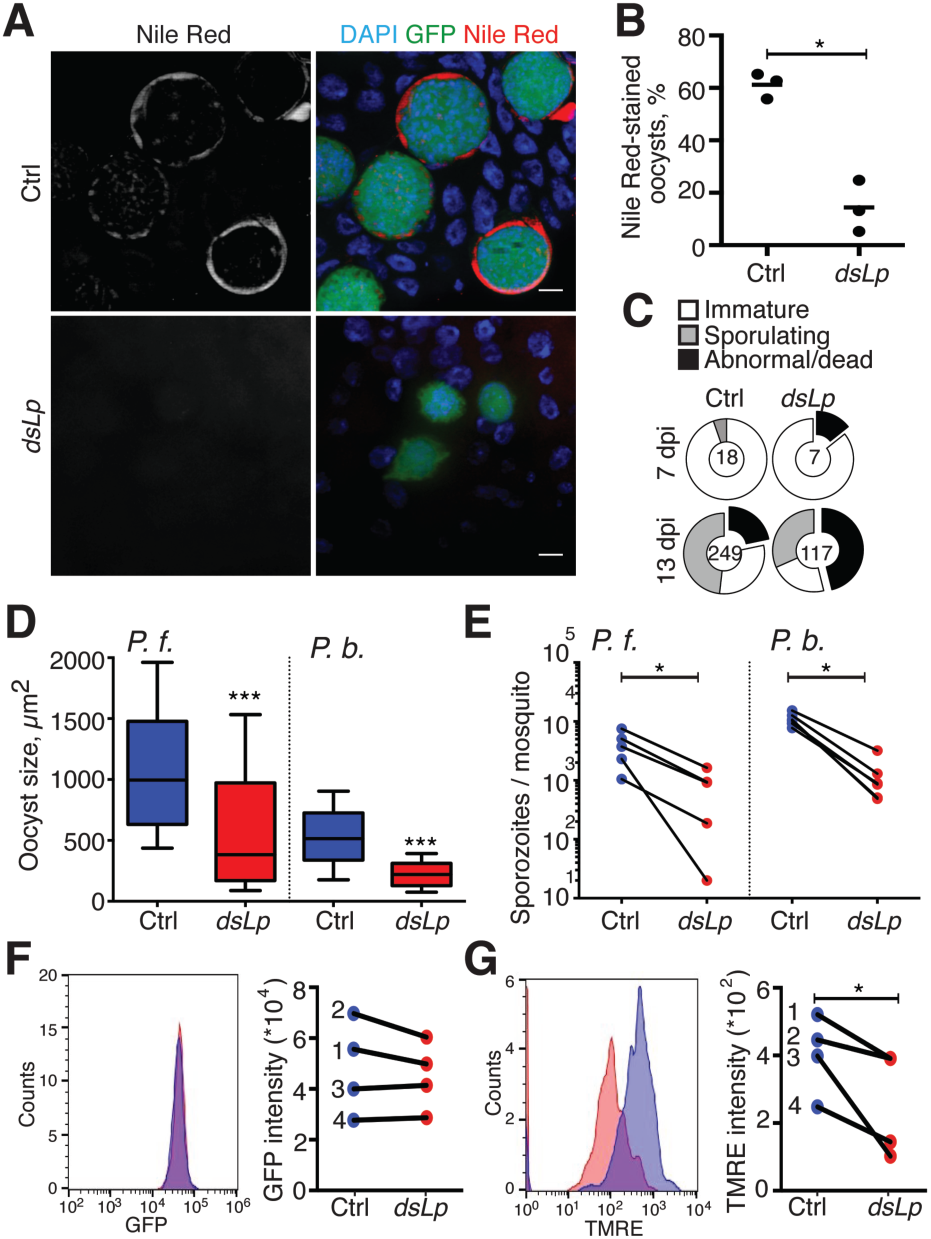
Mosquito lipids control within-vector oocyst proliferation, sporozoite metabolic activity and infectivity. Control and Lp-deficient female mosquitoes were infected with *P*. *berghei* (*P.b.*) (**A-G**) or *P. falciparum* (*P.f.)* (**D**, **E**) parasites. (**A**) Neutral lipids detected by the Nile Red stain in the GFP-expressing *P. berghei* oocysts (green) 14 dpi in control (Ctrl) and Lp-depleted (*dsLp*) mosquitoes. Nuclei are visualized by DAPI (blue). Scale bars −10 µm. (**B**) Percentage of Nile Red-positive *P. berghei* oocysts in control and Lp-depleted mosquitoes 13 dpi. At least 140 oocysts from 6 mosquitoes were analysed per condition. Each point represents mean of one experiment (n=3, *: p<0.05; paired t test). (**C**) Oocysts detected by transmission electron microscopy (see also Figure S3) were scored according to the developmental stage: non-sporulating oocysts with normal morphology (immature, white); oocysts with normal plasma membrane retraction, budding sporoblasts, or formed sporozoites (sporulating, gray); oocysts with aberrant electron-light intracellular vacuolization (abnormal/dead, black). Proportions of the categories are shown as pie charts and the total number of analyzed oocysts per condition is indicated in the middle of each chart. (**D**) Differences in oocyst sizes of *P.f.* (11 dpi, n=3) and *P.b.* (14 dpi, n=6) in control and Lp-depleted mosquitoes. At least 225 oocysts were measured per condition. The box plots represent medians (horizontal bars) with 10th and 90th percentiles. (***: p<0.0001, Mann-Whitney test). (**E**) Development of *P.f.* (14 dpi, n=5) and *P.b.* (18 dpi, n=4) salivary gland sporozoites in control and Lp-depleted mosquitoes. Each point represents the mean number of sporozoites per mosquito for each experimental replicate. Asterisks indicate statistically significant differences (*: p<0.05; 2-way ANOVA). Sporozoite (**F**) GFP expression and (**G**) mitochondrial membrane potential as measured by TMRE were analysed using imaging flow cytometry. Histograms from one representative experiment (left panels) and geometrical mean values from four independent experiments (right panels) in control (blue, n=1,405) and lipid-deprived (red, n=593) sporozoites are shown. Asterisks indicate statistically significant differences (*: p<0.05; paired t test).

As lipid restriction attenuated within-host virulence, we set out to identify *Plasmodium* traits affected by Lp depletion. In these experiments, we compared morphology, vitality and gene expression of the sporozoites produced by control and Lp-depleted mosquitoes. We found that lipid-deprived sporozoites were alive and properly expressed major maturation markers (Figure 3F and Figure S4). As lipids are crucial for metabolic activity, we measured mitochondrial membrane potential using TMRE dye, which accumulates in the membrane matrix of active mitochondria^23^. Lipid-deprived sporozoites showed a significant decrease in mitochondrial membrane potential compared to controls (Figure 3G and Figure S4), suggesting that mosquito lipids shape metabolic activity of *Plasmodium* transmissible forms. Moreover, TMRE intensities in control sporozoites were broadly distributed over a range of 2.5 logs, as opposed to a sharp peak of the cytoplasmic GFP signal (Figure 3F and G). These results suggest that genetically identical sporozoites display high variability in mitochondrial activity, which may contribute to the plasticity of within-host virulence. Taken together, our data provide experimental evidence that *Plasmodium* exploits Lp to scavenge mosquito lipid resources for its within-vector proliferation in a non-competitive parasitic manner, and identify mitochondrial membrane potential as a virulence-associated trait.

### Time-shift in *Plasmodium* sporulation circumvents competition for mosquito resources

What mechanism allows the parasites to avoid competition for the vector’s resource investment in reproduction? To answer this question, we gauged the timing of lipid consumption by the mosquito ovaries and of parasite proliferation. We observed a clear temporal shift between these two processes. While mosquito ovaries accumulated lipids within the first two days after feeding, *Plasmodium* proliferation began one week post-infection (Figure 4A). We hypothesized that in order to avoid a direct competition for nutrients, *Plasmodium* could have evolved to delay its replication until after completion of vector’s oogenesis. This time-shift hypothesis is compatible with the non-competitive parasitic scenario where parasites benefit from lipid delivery after completion of oogenesis, as long as sufficient resources are left for parasite development. Therefore, if lipid restriction affected only replicating oocysts, providing lipids at the initiation of oocyst sporogony should bypass one week of lipid starvation and rescue parasite development and virulence. We tested this hypothesis by directly gauging sporozoite infectivity and development into extra-erythrocytic forms (EEFs) in the hepatoma HepG2 cells *in vitro*.

**Figure 4.**
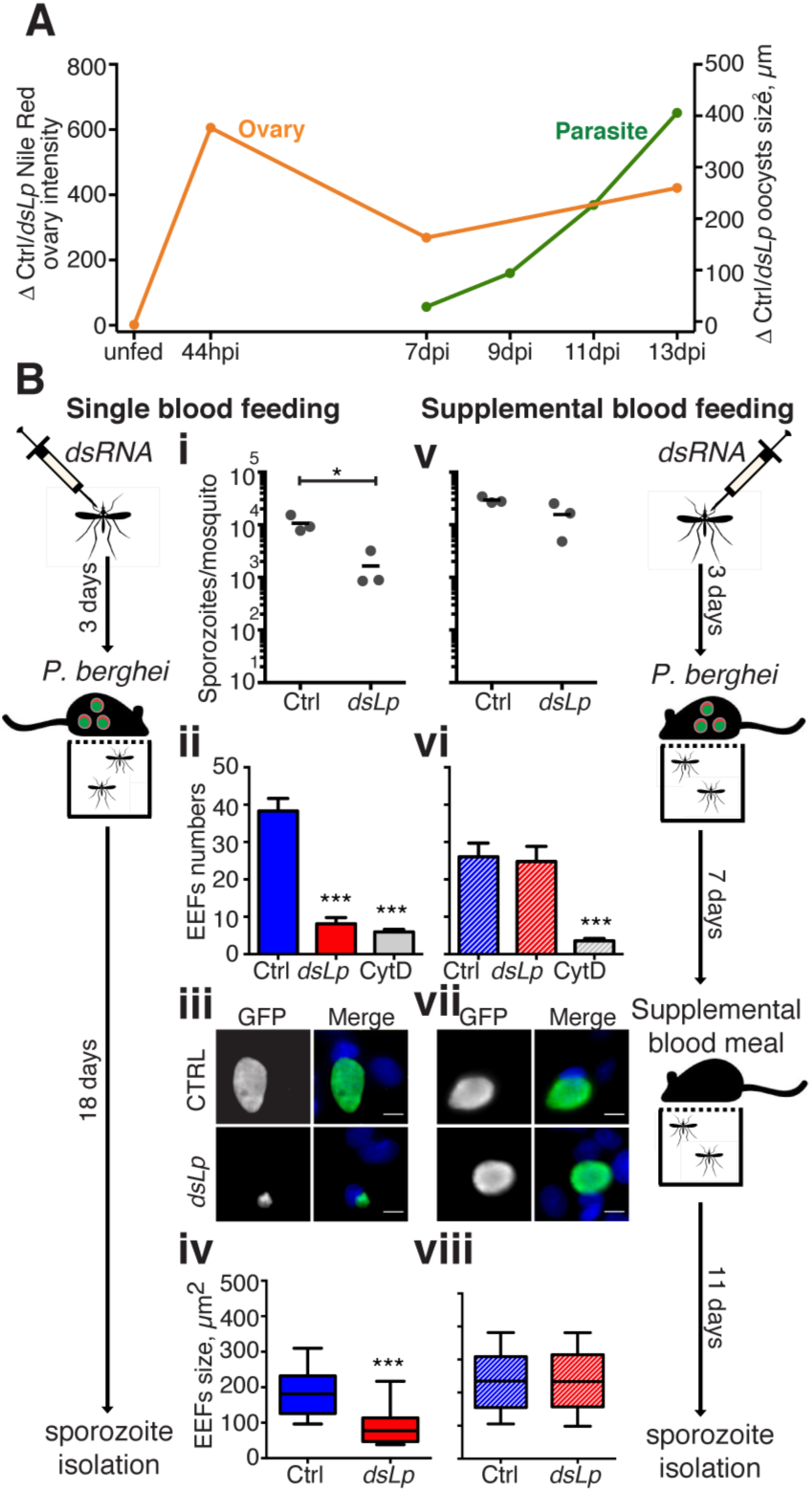
*Plasmodium* time-shift in sporulation circumvents competition for mosquito resources. (**A**) Kinetics of lipid acquisition by mosquito ovaries and *P. berghei* oocyst development. The ratios of Nile Red mean intensities in the ovaries (orange line) and oocysts growth (green line) between control and Lp-depleted mosquitoes are shown. Five or more mosquitoes were dissected per time point per condition (n≥2). (**B**) Lp-depleted mosquitoes (*dsLp*) and controls (Ctrl) were infected with *P. berghei* and offered (**v-viii**) or not (**i-iv**) a supplemental blood meal 7 days post infection. Salivary gland sporozoites were isolated and counted from control and Lp-deficient mosquitoes (**i** and **v**). Human hepatoma HepG2 cells were infected with 10,000 sporozoites and the number of extra-erythrocytic forms (EEFs) per field was gauged by microscopy 2 dpi (**ii** and **vi**). As a negative control, sporozoites from control mosquitoes were treated with cytochalasin D (Cyt D). Mean numbers and SEM are shown. Representative fluorescent micrographs of GFP-expressing EEFs (green) developing in HepG2 cells (**iii** and **vii**). Cell nuclei are stained with DAPI (blue). Scale bar - 10 µm. EEF sizes produced by sporozoites from control and Lp-deficient mosquitoes (**iv** and **viii**). The box plots represent 10th and 90th percentiles while the horizontal lines show medians. Asterisks indicate statistically significant differences (n=3, *: p<0.05; ***: p<0.0001; Kruskal-Wallis and Mann-Whitney test).

Cells were exposed to equal numbers of sporozoites isolated from control and Lp-depleted mosquitoes that received (Figure 4Bv-viii) or not (Figure 4Bi-iv) a supplemental blood feeding seven days post infection, at the time when the first effects of lipid starvation on the oocyst’s size were observed (Figure 3C and Figure S3). EEF development was examined by fluorescence microscopy. As expected for a single blood feeding, lipid restriction attenuated sporozoite numbers, infectivity and development *in vitro*, as significantly smaller EEFs were produced by lipid-deprived sporozoites compared to controls (Figure 4Bi-iv and Figure S5). Strikingly, the supplemental feeding restored sporozoite numbers in Lp-depleted mosquitoes, as well as parasite infectivity and EEF development (Figure 4Bv-viii and Figure S5). Our data show that delivering lipids to the oocysts after completion of vector’s oogenesis rescues *Plasmodium* infectivity and virulence. Taken together, these results suggest that a time-shift in *Plasmodium* sporogony circumvents within-vector competition for reproductive resources.

## DISCUSSION

Virulence is one of the most evolutionarily relevant and complex pathogen traits. Understanding virulence mechanisms provides an opportunity to predict and, potentially control the pathogen burden and its evolution. Here we generated theoretical models of the mosquito-parasite relationship, experimentally evaluated them, and found that *Plasmodium* hijacks the main mosquito lipid transporter for lipid delivery to the actively proliferating oocysts after completion of the vector reproductive cycle. Such time-shift in oocyst maturation benefits the parasitic relationship by allowing a timely allocation of resources needed for host reproduction and for parasite proliferation. Temporal regulation plays a critical role in host-parasite interactions and in host immune responses. In mosquitoes, the ookinetes at the basal side of the midgut are the major target of the mosquito immune system within the first 24-48 h after blood meal, whereas even the earliest oocyst stages are spared from the complement attack^24^. Interestingly, sporogony is the longest replicative process in the life cycle of all *Plasmodium* species (12 to 16 days depending on the species). Given the average mosquito life span of 30 days, it roughly corresponds to a half of mosquito life. Prolongation of the within-vector sejour is risky for the parasite, which should be selected to maximize its chances to infect the next host as fast as possible. Therefore, the beneficial effects of lengthy sporogony must be under a strong selective pressure. We predict that shifting the timing of resource allocation should have dramatic consequences for parasite development, transmission and virulence.

Our observations suggest that limited lipid resources during sporulation induce an autophagy-like cell death and select for the most competitive parasites that effectively scavenge nutrients from the mosquito. Autophagy, often characterized by cytoplasmic vacuolization, has been observed in a large proportion of *Plasmodium* liver stages^25^, and our results extend these observations to the extracellular oocysts. Nutrient restriction in natural conditions could be induced by oocyst crowding^19^; however in our experiments the numbers of pre-sporogonic oocysts between the experimental groups were similar^21^. Interestingly, prolonged periods of nutrient deprivation cause cell autophagy and induce mitochondrial dysfunction in mouse embryonic fibroblasts^26^. Similarly, we found that lipid restriction blunted sporozoite’s mitochondrial membrane potential. Mitochondria are essential for mosquito stages of the *Plasmodium* life cycle, as mutations in mitochondrial genes arrest growth and sporulation of oocysts^27–30^. Moreover, inhibition of mitochondrial potential by the antimalarial drug atovaquone inhibits *Plasmodium* sporogonic development and reduces the number and size of developing EEFs *in vitro* ^31–33^. Together with these findings, our results identify mitochondria as crucial regulators of within-vector *Plasmodium* proliferation, transmission and within-host virulence. Although drug pressure on the mitochondria frequently selects for resistance, most of the identified mutations are loss-of-function, that disrupt within-vector parasite development and disable transmission^30^. Therefore, *Plasmodium* mitochondria may represent an evolution-proof drug target^34^, as long as no gain-of-function mutations emerge that may increase mitochondrial activity and, thereby, parasite virulence.

At the ecological level, our data show that the plasticity of parasite virulence is metabolically regulated. We found that even in normal conditions, sporozoites display broad variability in their mitochondrial activity. This implies that mosquito life-history traits can directly influence parasite virulence. Variable trophic environments of larval breeding sites^35^ and of blood meals should translate inter-individual mosquito nutritional differences into variability in *Plasmodium* virulence. Indeed, multiple reports have implicated larval diet and adult blood meals in determining parasite loads within a mosquito^18,36–40^, but this has not been linked to within-vector modulation of parasite metabolic activity and within-host virulence. Our findings open up a new perspective on mosquito transmission capacity and on evolution of malaria virulence.

The evolutionary responses to vector control measures are often non-linear due to complexity of vector-parasite relationships. Competitive within-vector *Plasmodium* dynamics may result in a virulence runaway, leading to a competitive exclusion between vector and parasite and exacerbating within-host virulence. We show that parasitic non-competitive exploitation of vector lipid resources restricts evolution of *Plasmodium* virulence. Importantly, harm towards the vector’s reproduction alone does not restrict virulence evolution^41^. The trade-off between within-vector parasite density and within-host virulence that restrains virulence at intermediate levels, emerges only when parasite resource acquisition depends on vector mechanisms. We provide empirical evidence that within-vector *Plasmodium* development relies on the major vector lipid transporter and that competition between vector and parasite is prevented by a time-shift between vector oogenesis and parasite sporogony. This beneficial developmental niche should be retained as long as nutrient supply is sufficient. In a low nutrient environment, the relationship may switch towards competition, which would rapidly invert the trajectory of virulence evolution. Further investigation of the effects of other mosquito factors on parasite resource acquisition is necessary to predict potential switches between parasitic and competitive vector-parasite relationships. Such situations could arise from targeting mosquito reproduction factors essential for parasite development by transgenic approaches^42^, or from introduction of such biological competitors as *Wolbachia* ^43^. All together our data highlight the links between metabolism and life-history traits of vector and parasite, that shape evolutionary trajectories of malaria virulence. Further theoretical, experimental, and field studies are needed to determine the effects of natural and artificial perturbations of vector and parasite metabolism on malaria epidemiology.

## MATERIALS AND METHODS

### Mathematical modeling

#### Within-vector dynamics

Vector and parasite compete for resources from a single blood meal (non-replenished nutrient source). To investigate how vector-parasite relationship shapes within-host virulence, we first modeled resource competition between vector reproduction and sporozoite development. For this purpose, we use a Lottka-Volterra competition model^14,15^, considering inter-and intraspecific competition between mosquito eggs (*E*) and parasites (*P*):

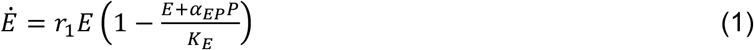

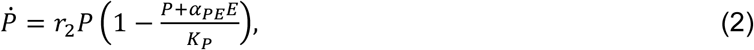

where *r*_1_ and *r*_2_ represent the maximum intrinsic growth rate of eggs and parasites, respectively, and *α*_*EP*_ and *α*_*PE*_ describe their competition strength. The maximum egg and sporozoite densities are given by the carrying capacities *K*_*E*_ and *K*_*P*_ which from this point on we assume to be 1.

The four equilibrium points are given by

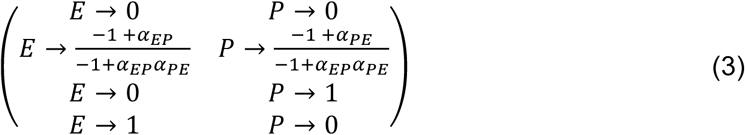

and describe three qualitatively different scenarios: extinction of both species, competitive exclusion, or co-existence. Because both eggs and parasites are found in the mosquito, we hereafter only focus on the non-trivial steady state, i.e., co-existence, which can only occur if both growth rates (*r*_1_ and *r*_2_) are positive and neither species outcompetes the other, i.e., if *α*_*EP*_ and *α*_*PE*_ are smaller than 1.

For all cases, where 0 < *α*_*EP*_ < 1; *and* 0 < *α*_*PE*_ < 1, sporozoite production 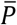 decreases with increasing values of *α*_*EP*_ and increases with increasing values of *α*_*EP*_ for all, where 0 < *α*_*EP*_ < 1; 0 > *α*_*PE*_ < 1 (Figure S6). Instead, vector reproduction Ē should decrease with increasing levels of *α*_*EP*_ independent of the sign of *α*_*PE*_ (Figure S7).

#### Epidemiological dynamics

Following Alizon and van Baalen^16^, *Plasmodium* epidemiological reproductive ratio R_0_ is a product of vector-to-host 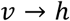 and host-to-vector 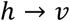 transmission rates *β*, and of the availability of susceptible hosts *S*. These are constrained by the respective duration of infection defined by parasite induced mortality 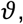, recovery rate *γ*, and intrinsic host mortality *µ*:

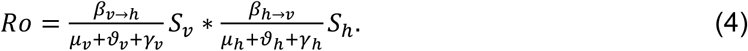

#### Nested model

To integrate the within-vector dynamics into an epidemiological model of malaria transmission, we first assumed that within-vector sporozoite production is at equilibrium before the time of transmission (i.e., quasi-steady state). Because vector-to-human transmission rate increases with the number of produced sporozoite^44^, we next postulated that vector-to-human transmission rate is a linear function of equilibrium parasite density 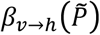:

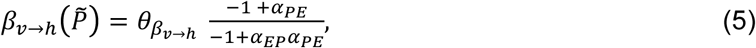

where 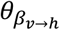 is a constant which describes the maximum transmission rate in relation to parasite density.

Host-to-vector transmission depends on the parasites asexual replication and production of sexual forms (gametocytes)^5,45^. Virulence towards the host is caused by the asexual production during the blood stages, which makes host-to-vector transmission a linear function of virulence 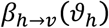. We assume that the capacity to enter the asexual cycle depends on initial parasite load, which links parasite density at the time of vector to human transmission to human to vector transmission:

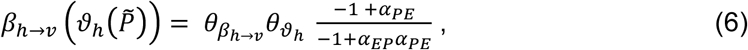

where 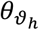 and 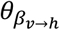 are the rates for virulence and transmission in relation to parasite loads. However, within-vector resource competition may also qualitatively affect the parasite’s capacity to infect the human host^17-19^. Therefore, we assumed that within-host virulence increases with the efficiency of resource exploitation via the parasite’s competition coefficient:

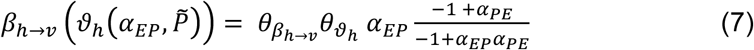

R_0_ under these assumptions is given by equation 8. It can be used to study the trajectories of virulence evolution in diverse vector-parasite relationships. The parameter values used in this study are provided in Table S1.

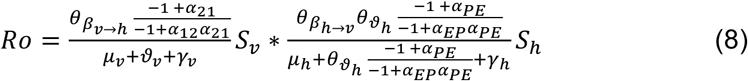

### Mosquito rearing and parasite Infections

*Anopheles gambiae s.l.* mosquitoes were used throughout the study. To prevent large losses at early parasite stages (ookinetes) inflicted by the mosquito complement-like system in the absence of Lp^21,46^, all infection experiments were performed with immune-deficient mosquitoes (*7b* line)^47^. G3 and *7b* mosquitoes, were maintained at 29°C 70%– 80% humidity 12/12 h day/night cycle. In *P. falciparum* infections, mosquitoes were fed at 37°C for 15 min on a membrane feeder with NF54 gametocytes cultured with O+ human red blood cells (Haema, Berlin), and, thereafter, kept in a secured S3 laboratory according to the national regulations (Landesamt für Gesundheit und Soziales, project number 411/08). For *P. berghei* experiments, mosquitoes were fed on anesthetized CD1 mice infected with the GFP-expressing *P. berghei* GFP-con 259cl2 clone (ANKA strain)^48^. Shortly after infections, unfed mosquitoes were removed, while fed mosquitoes were maintained at 26°C for 11-14 days (*P. falciparum*) or at 20°C for 7-21 days (*P. berghei*), and then used for the midgut and/or the salivary gland dissections.

### RNAi silencing

Double stranded RNA (dsRNA) against *lipophorin* (*dsLp*) was produced as previously described^21^. For RNAi silencing, 1-2 day-old females were anesthetized with CO_2_ and injected with 69 nl of 3 µg/µl *dsLacZ* (control) or *dsLp* using a Nanoject II Injector (Drummond). Mosquitoes recovered for 3-4 days following injection before infections. Efficiencies of RNAi silencing are summarized in Figure S8.

### Immunofluorescence analysis

Mosquito midguts were dissected in PBS, fixed in 4% formaldehyde and washed in PBS. Tissues were permeabilized with 0.1% Triton X-100 and incubated with a 1:1 mix of mouse monoclonal anti-ApoI and anti-ApoII antibodies (1/300, clones 2H5 and 2C6)^21^ overnight at 4°C followed by incubation for 40 min at room temperature with the secondary Cy3-labeled antibodies at 1/1,000 (Molecular Probes), or with 0.1 µg/ml of the Nile Red stain (Sigma Aldrich). Nuclei were stained with DAPI (1.25 µg/ml, Molecular Probes) for 40-60 min at room temperature. Images were acquired using an AxioObserver Z1 fluorescence microscope equipped with an Apotome module (Zeiss). Oocyst size was measured by hand-designing a circular ROI of randomly selected oocysts (identified by GFP for *P. berghei* or by bright field and DAPI staining for *P. falciparum*) using ZEN 2012 software (Zeiss). Images were then processed by FIJI software (ImageJ 1.47m).

### Development of *P. berghei* liver-stages *in vitro*

The salivary glands of *P. berghei* infected mosquitoes were dissected, sporozoites were collected into RPMI medium (Gibco) containing 3% bovine serum albumin (Sigma Aldrich) and enumerated with a hemocytometer. The *P. berghei* liver stages were cultured *in vitro* in HepG2 hepatoma cells and analyzed using standard techniques^49^. Briefly, 15,000 −20,000 HepG2 cells per well (70% confluence) were plated on transparent-bottom 96-wells plates (Nalgene International) and incubated for 24 h, then seeded with 10,000 sporozoites and co-cultured at 37°C for 2 h. Free sporozoites were removed by washing. As a negative control, sporozoites isolated from control mosquitoes were pre-treated with the inhibitor of actin polymerization Cytochalasin D (Sigma Aldrich) for 10 min. At 48 h post seeding, the cells were fixed with 4% formaldehyde for 10 min, washed with PBS and blocked with 10% fetal calf serum in PBS. Development of the liver-stage parasites was examined using monoclonal mouse anti-GFP antibodies (1/1,000, AbCam) and revealed with the secondary AlexaFluor488 conjugated anti-mouse antibody (1/1,000, Molecular Probes). Nuclei were stained with DAPI (1.25 µg/ml, Molecular Probes). Images were recorded directly on the 96-well plate using an AxioObserver Z1 fluorescence microscope equipped with an Apotome module (Zeiss) and analyzed for number and size of liver forms using the Axio-Vision ZEN 2012 software (Zeiss).

### *P. berghei* mouse infections *in vivo*

Mice were housed and handled in accordance with the German Animal Protection Law (§8 Tierschutzgesetz) and both institutional (Max Planck Society) and national regulations (Landesamt für Gesundheit und Soziales, registration number H 0027/12). To determine infectivity to mice, the sporozoites collected from the mosquito salivary glands were injected subcutaneously (5,000 sporozoites/mouse) into the tails of 8-10-week-old C57BL/6 females. Bite-back experiments were performed by feeding the *P. berghei*-infected mosquitoes (18 dpi) on anesthetized naïve C57BL/6 mice (mean 9 bites/mouse, range 2-15). Parasitemia was determined by daily Giemsa staining of thin blood smears and FACS analysis of the red blood cells with the GFP-expressing blood stage parasites. Infected mice were monitored every 6-12 h for the appearance of severe neurological and behavioral symptoms typical of experimental cerebral malaria (ECM) such as hunched body position, grooming alteration, ataxia, paralysis, or convulsions^50^ or by the rapid neurological and behavioral test (RMCBS)^51^. All mice with ECM symptoms or the RMCBS score equal or below 5/20 were sacrificed immediately.

### Sporozoite imaging flow cytometry

Salivary gland sporozoites were isolated into RPMI medium (Gibco) 3% bovine serum albumin (Sigma Aldrich) and kept on ice until staining. Sporozoites were diluted in PBS to concentrations corresponding to 3 mosquito equivalents and stained in the dark with TMRE (5 nM) (Cell Signaling Technology) for 20 min at 20°C. Images were acquired without washing using an ImageStreamX Mk II (Merck Millipore) with a 60x objective over 1 h period. To avoid a possible bias due to the variable pre-acquisition waiting times on ice, the order of sample acquisition was swapped between the experimental replicates. GFP-expressing sporozoites were gated by size (Area_M01 or Area_M02) and by GFP intensity (Intensity_MC_Ch02). The analysis was performed with IDEAS 6.2 (Merck Millipore) and FCS files were exported and analyzed by FlowJo v10. Given the small size of the mitochondria, the feature finder tool from IDEAS software was used to improve gating of optimally focused sporozoites. The following parameters were used to discriminate in-focus single sporozoites from debris, and to identify in-focus sporozoites to resolve their mitochondria: H Entropy Std_M01_Ch01_15 over H Variance Std_M01_Ch01, Aspect Ratio_M02 over Symmetry 4-Object(M02_Ch2) Ch02, Gradient RMS-M01-Ch01 over H Correlation Mean-M02-Ch02_3, and Gradient RMS-M02_Ch02. Briefly, these parameters measure brightfield image texture, GFP image shape, brightfield and GFP focus, respectively. For quantification of mitochondrial membrane potential and GFP intensity, each sample was time-gated to exclude artefacts of prolonged staining. The parameter corresponding to the lowest background signal in unstained sporozoites was Bright Detail Intensity R7_M04_Ch04, Geo Mean which was selected as *bona fide* TMRE intensity readout (see Figure S5D). For GFP intensity, the Intensity_MC_Ch02, Geo. Mean was used. To validate the use of TMRE as a marker for sporozoite membrane potential, we applied 50 µM of a mitochondrial uncoupling drug CCCP (Cell Signaling Technology) and showed that drug treatment abolished TMRE signal in sporozoites (see Figures S4D).

### Statistical analysis

Statistical analysis was performed with GraphPad Prism 7 software and p values below 0.05 were considered significant (*: p<0.05; **: p<0.001; ***: p<0.0001) and indicated in the figures. The specific tests used are indicated for each figure in the corresponding legend.

## ACKNOWLEDGEMENTS

This work was supported by Max Planck Society, EC FP7 EVIMalaR (grant agreement n°242095) and MALVECBLOK (grant agreement nº223601). The authors thank H. Krüger, M. Andres and L. Spohr for mosquito rearing, mouse work assistance, and *Plasmodium* infections, and D. Tschierske and D. Eyermann for *P. falciparum* cultures. The authors express their gratitude to Prof. K. Matuschewski and his group for continuous fruitful discussions, to Dr. M.M. Mota and Dr. P. Carrillo-Bustamante for helpful comments.

## AUTHOR CONTRIBUTIONS

GC and EAL conceived the study and designed the experiments. MG performed the modeling analysis. GC, ME, RL, CG, and VB performed experiments. AEH and RS contributed reagents and expertise. GC, ME, MG, RL, RS, CG, VB, and EAL analyzed the data. GC, MG and EAL wrote the manuscript.

